# Spatial sorting enables comprehensive characterization of liver zonation

**DOI:** 10.1101/529784

**Authors:** Shani Ben-Moshe, Yonatan Shapira, Andreas E. Moor, Keren Bahar Halpern, Shalev Itzkovitz

## Abstract

The mammalian liver is composed of repeating hexagonal units termed lobules. Spatially-resolved single-cell transcriptomics revealed that about half of hepatocyte genes are differentially expressed across the lobule. Technical limitations impede reconstructing similar global spatial maps of other hepatocyte features. Here, we used zonated surface markers to sort hepatocytes from defined lobule zones with high spatial resolution. We applied transcriptomics, microRNA array measurements and Mass-spectrometry proteomics to reconstruct spatial atlases of multiple zonated hepatocyte features. We found that protein zonation largely overlapped mRNA zonation. We identified zonation of key microRNAs such as miR-122, and inverse zonation of microRNAs and their hepatocyte gene targets, implying potential regulation through zonated mRNA degradation. These targets included the pericentral Wnt receptors Fzd7 and Fzd8 and the periportal Wnt inhibitors Tcf7l1 and Ctnnbip1. Our approach facilitates reconstruction of spatial atlases of multiple cellular features in the liver and in other structured tissues.

## Introduction

The mammalian liver is a highly structured organ, consisting of repeating hexagonally shaped units termed ‘lobules’ (Fig. 1a). In mice, each lobule consists of around 9-12 concentric layers of hepatocytes^1^. Blood flowing from portal nodes (PN) at the corner of the lobules towards draining central veins (CV) generates gradients of oxygen, nutrients and hormones along the lobule radial axis. Additionally, Wnt morphogens secreted by endothelial cells surrounding the CV create a graded morphogenetic field^2^. This graded microenvironment gives rise to spatial heterogeneity in gene expression among hepatocytes residing at different lobule layers, a phenomenon that has been termed ‘liver zonation’^3,4^.

**Fig 1.**
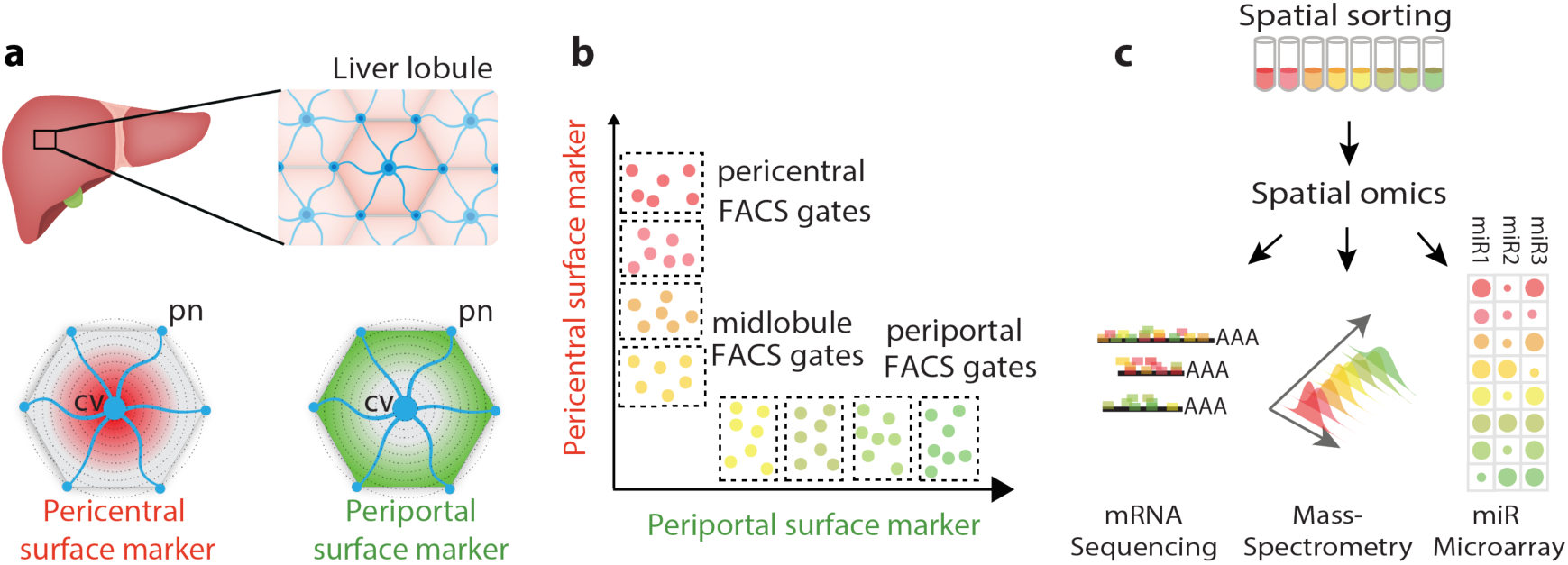
Spatial sorting approach for isolating large amounts of hepatocytes from distinct layers with high resolution. **a**, Identification of zonated surface markers. cv – central vein. pn – portal node. **b**, Fluorescence-activated cell sorting (FACS) enables defining gates that enrich for zonated hepatocytes according to their surface marker expression. **c**, Spatially-sorted hepatocytes can be measured using multiple assays that require large input material, such as RNA-seq, Mass spectrometry and microRNA microarray applied in the current study.

We have recently used spatially-resolved single cell transcriptomics to uncover the global zonation patterns of hepatocyte gene expression^5^. We found that around half of all genes expressed in hepatocytes are zonated, with specific functional specialization that seem to match the zonated microenvironment. This global zonation suggests that similar spatial heterogeneity of hepatocytes may also exist for other cellular features, including proteins, metabolites and regulatory molecules such as microRNAs. However, achieving similar global zonation maps for cellular features beyond mRNA has encountered technical difficulties.

Immunohistochemistry enables measurements of protein levels with high spatial resolution but it is low-throughput and often limited by lack of availability of antibodies. Laser capture microdissection (LCM) and digitonin perfusion enable extracting large numbers of periportally or pericentrally enriched cells^6,7^. However, these techniques are limited in spatial resolution and purity, since they measure both hepatocytes and non-parenchymal cells. Single cell measurements of cellular features beyond mRNA are starting to emerge^8,9^, however these technologies are less mature in tissues, compared to bulk analysis. A methodology that would enable massive isolation of pure cell types from defined layers with high spatial resolution would enable generating organ spatial atlases of key features such as methylation patterns, chromatin conformations, microRNA content and proteomics. In the liver, such measurements would broaden our understanding of the regulation of liver zonation and could be used in order to more precisely model liver metabolic function.

In this work, we developed an approach termed ‘spatial sorting’, that utilizes surface markers with discordant zonation profiles to isolate massive amounts of hepatocytes from defined lobule layers (Fig. 1b). We used these for high-throughput profiling of mRNAs, microRNAs and proteins (Fig. 1c), revealing novel features of liver zonation. These include a comprehensive proteomic zonation map and the identification of zonated microRNA with discordantly zonated target genes. Our approach can be readily applied to profile other cellular features of hepatocytes and other cell types in health and disease.

## Results

### Spatial sorting enables isolating bulk hepatocyte populations from different lobule layers with high spatial resolution

We used our recently reconstructed mRNA zonation map^5^ to identify zonated surface markers with a large dynamic range in expression, spanning several radial lobule layers (Fig. 1a, Supplementary Fig. 1a). We argued that the combined staining of two inversely zonated surface proteins would be informative for inferring the lobule positions of single hepatocytes (Fig. 1b), which would facilitate cell sorting of many cells according to their spatial origin (Fig. 1b-c). CD73, encoded by the gene Nt5e, is an enzyme converting mononucleotides to nucleosides that exhibits pericentral zonation. E-cadherin, a cell-cell adhesion glycoprotein encoded by Cdh1, exhibits periportal zonation^10^ (Fig. 2a). We used immunofluorescence to validate the zonation of these two surface markers at the protein level as well (Fig. 2b-c).

**Fig 2.**
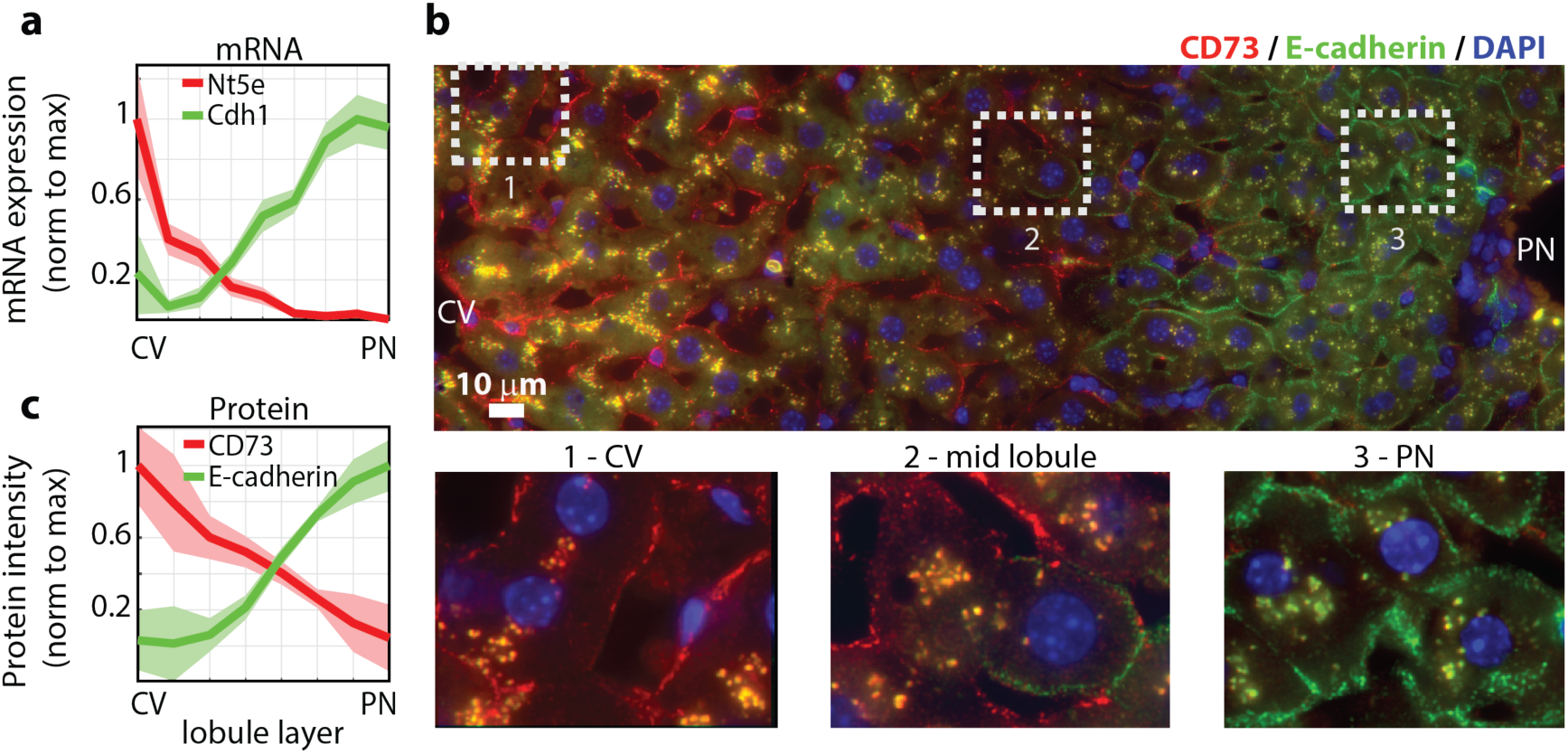
CD73 and E-cadherin are inversely zonated surface markers. **a**, Nt5e, encoding CD73 and Cdh1, encoding E-cadherin are surface markers that are zonated at the mRNA level. Data taken from ^5^. **b**, CD73 and E-cadherin proteins are zonated. Shown is an example of a lobule stained by immunofluorescence with antibodies against CD73 (red) and E-cadherin (green). Cytoplasmic yellow blobs are tissue auto-fluorescence. Blue – DAPI nuclear stain. **c**, Quantification of immunofluorescence images (8 lobules from three mice). Patches are SEM across the eight lobules. CV – central vein, PN – portal node.

We perfused livers to dissociate single cells and performed Fluorescence-activated cell sorting (FACS) of isolated hepatocytes stained with antibodies against CD73 labeled with APC and E-cadherin labeled with PE (Methods). We filtered hepatocytes by size and by selecting cells that were negative for the endothelial cell marker CD31 and the immune cell marker CD45, to avoid pairs of hepatocytes and non-parenchymal cells (NPCs)^11^. We further filtered out non-viable cells and selected 4n hepatocytes using Hoechst staining (Fig. 3a). Stratifying hepatocytes by ploidy was important to obtain precise lobule localization (Methods, Supplementary Fig. 1b-c). The selected hepatocytes displayed strong anti-correlation in the fluorescence of CD73 and E-cadherin, as expected from the zonated expression patterns (Fig. 3b).

**Fig 3.**
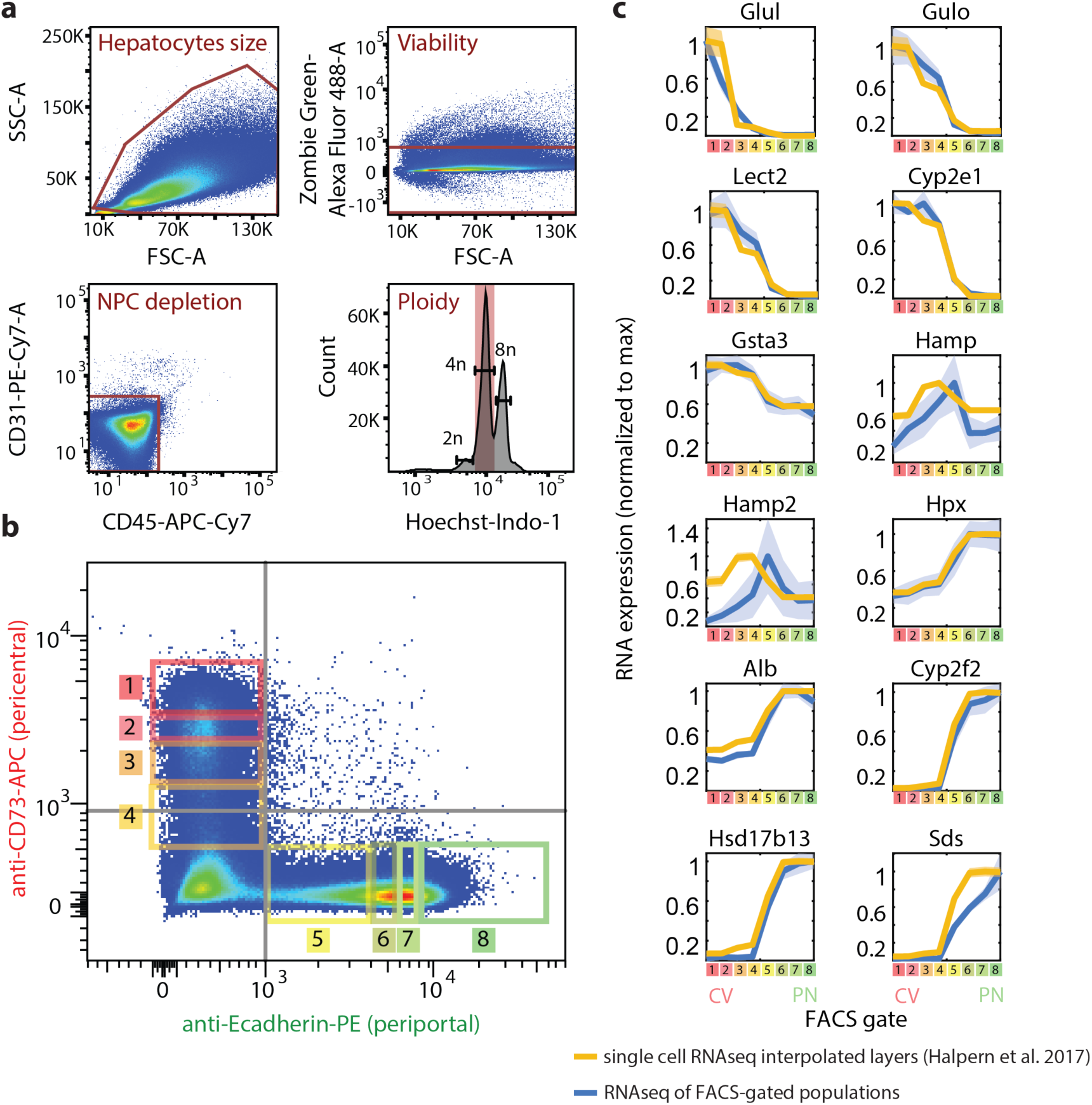
Spatial sorting reliably captures the different lobule layers. **a**, FACS gating strategy. FSC-A and SSC-A were used for hepatocytes size selection. Non-viable cells were filtered out by Zombie Green Viability kit. Staining with CD31 and CD45 antibodies enabled to gate out non-parenchymal cells. Tetraploid hepatocytes were selected based on Hoechst stain. **b**, Distribution of the included cells (40-60% from all events) according to intensities of CD73 and E-cadherin. Grey lines mark the unstained control limits, rectangles and numbers mark the gates used for spatially-sorted populations. **c**, Max-normalized expression patterns of selected genes along the different FACS gates in blue (N=5 mice), compared with interpolated max-normalized zonation profiles based on ref.5 (Methods) in yellow. Line patches represent SEM.

We defined eight gates based on the combined fluorescence of CD73 and E-cadherin (Fig. 3b). To ensure reproducibility, the gates were defined as percentiles of the marginal expression levels of each surface marker, compared to unstained control (Methods). To validate that our defined gates represent sequential lobule layers, we performed bulk RNA sequencing (RNAseq) on 10,000 sorted hepatocytes from each gate and compared the zonation profiles to our spatially-resolved scRNAseq map^5^. Zonation profiles were highly concordant (Fig. 3c, Supplementary Table 1), demonstrating the feasibility of our approach for isolating bulk hepatocytes with high spatial resolution.

### Mass spectrometry proteomic measurements of spatially-sorted hepatocytes

We next applied spatial sorting to reconstruct the zonation patterns of the hepatocyte proteome. To this end, we sorted 100,000 hepatocytes from each of the eight FACS gates for five *ad-lib*. mice and performed mass-spectrometry proteomics. For each mouse and gate we also isolated 10,000 cells and applied bulk RNAseq. The Mass-spectrometry measurements yielded 3,210 identified proteins (Methods, Supplementary Table 2). The hepatocyte protein content averaged over all FACS gates was highly correlated with a previous bulk measurements compared to ref.^12^, with Spearman’s r = 0.75, (Supplementary Fig. 2).

Our dataset included 3,051 proteins with matched mRNA (Supplementary Table 3). The means of the protein and mRNA levels over all gates were positively correlated (Spearman’s r = 0.5, p-val = 1.2×10^−181^). Yet, for some proteins, there was a marked difference in protein and mRNA relative abundances (Fig. 4). These predominantly included hepatocyte secreted proteins. For example, Alb, encoding the secreted carrier protein albumin, was the most highly abundant hepatocyte mRNA (0.050±0.004 of cellular transcripts) but was ranked only 64 in protein levels (0.0034±0.0002 of cellular proteins). Other secreted proteins, which were ranked significantly higher in mRNA compared to the protein level included apolipoproteins encoded by Apoa1, Apoa2, Apoe, alpha-antitrypsin encoded by Serpin genes, complement system proteins and vitronectin, encoded by Vtn (Fig. 4a). A similar discordance between the levels of mRNAs and proteins for secreted genes was previously observed in mammalian cell lines^13^. Ribosomal mRNAs and proteins had a protein to mRNA ratio close to one, whereas genes of the TCA cycle had substantially higher protein to mRNA levels (Fig. 4b, Supplementary Fig. 2). Cps1, encoding the urea cycle enzyme carbamoyl-phosphate synthase was ranked first in protein content (0.0682±0.0066), but only 478 in mRNA expression (2.88×10^−4^±5.4×10^−5^ of cellular transcripts, Fig. 4a). Thus, the relative expression levels of mRNAs and proteins differ for distinct functional classes.

**Fig 4.**
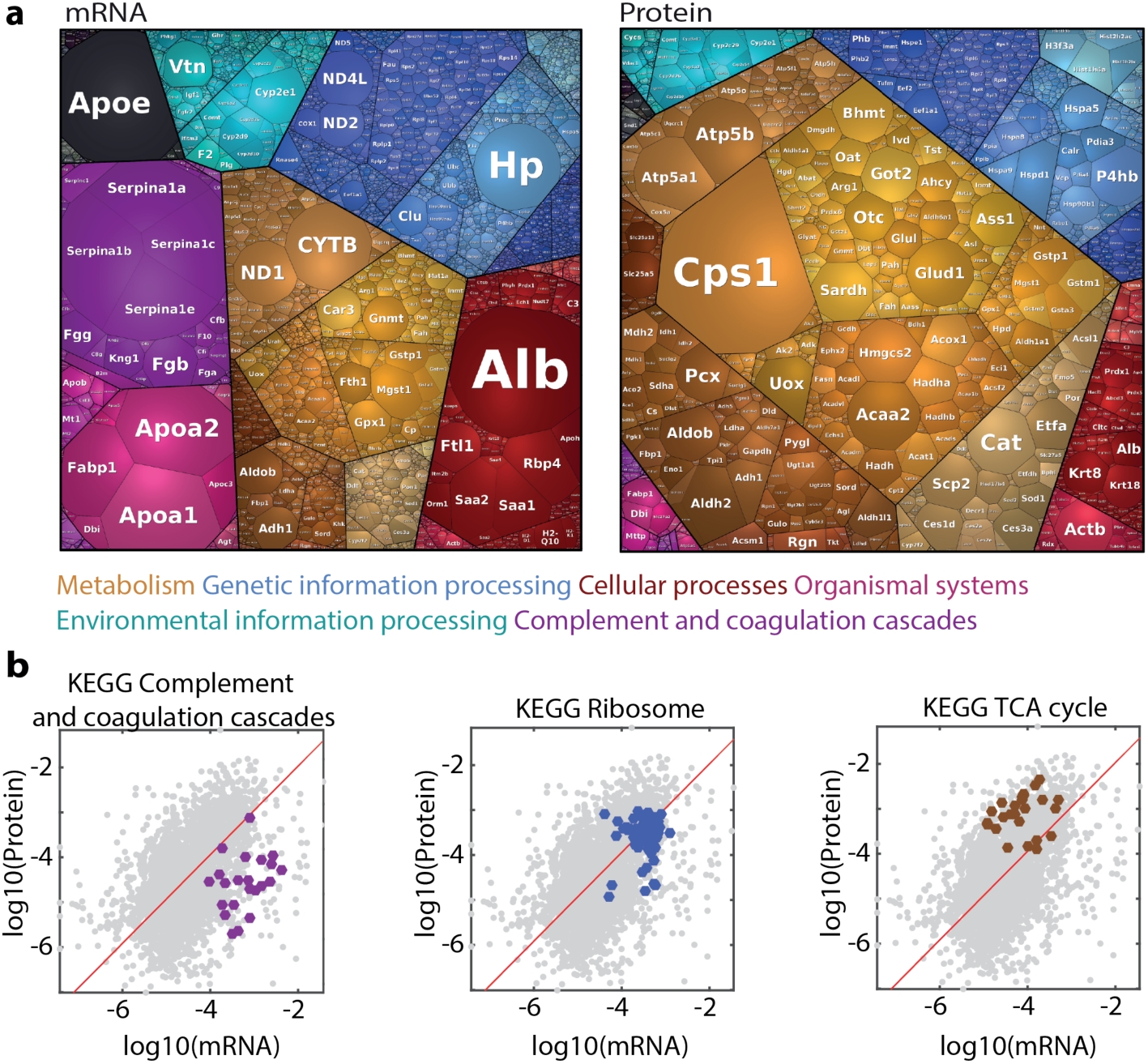
Correlations between mRNA and protein levels. **a**, Proteomaps for visualizing the distributions of the mean mRNA and mean proteins over all FACS gates. Each tile represents a gene, size is proportional to its fraction in the total dataset. Visualization was done using https://bionic-vis.biologie.uni-greifswald.de/^14–16^. Color classification key for selected categories is shown at the bottom. **b**, Gate-averaged mRNA and protein levels are mildly correlated (Spearman r=0.5). Shown are three KEGG functional classifications with distinct ratios of mRNAS and proteins. Red is a linear regression line (Methods).

### Zonation patterns of the hepatocyte proteome

We next examined whether the hepatocyte proteome exhibited zonated patterns. We found that 55% of the hepatocyte proteins (1,672 out of 3,051) were significantly zonated (FDR<0.05, Kruskal-Wallis test, Fig. 5a). Periportal and pericentral enriched KEGG pathways largely recapitulated previous zonation studies. Bile acid biosynthesis, lipid metabolism and P450 xenobiotic metabolism were pericentrally zonated, while gluconeogenesis, oxidative phosphorylation and complement and coagulation cascades were periportally zonated (Supplementary Fig. 3).

**Fig 5.**
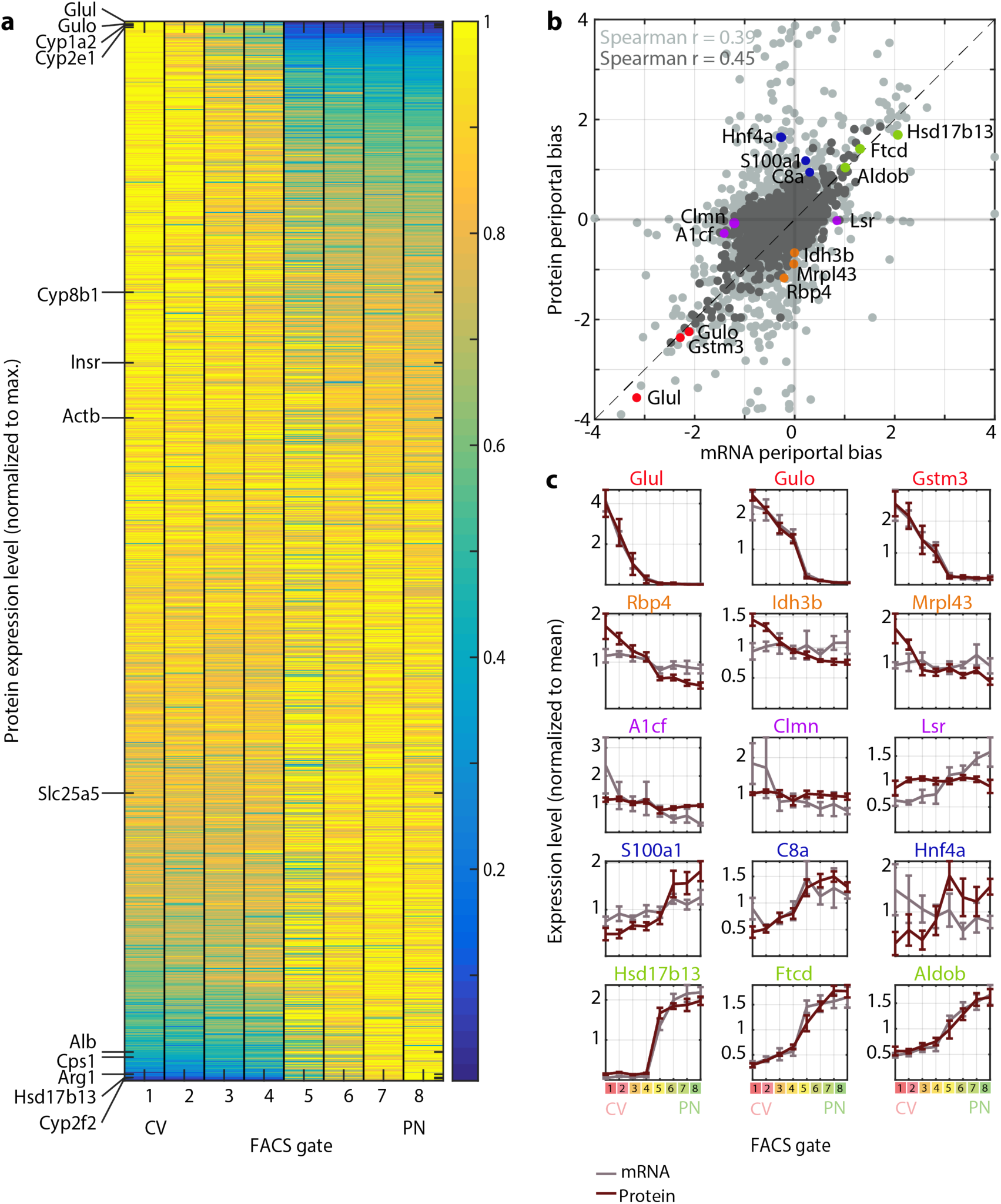
A spatial atlas of the hepatocyte proteome. **a**, Zonation of hepatocyte proteins. Genes are sorted by the zonation profile center of mass. Selected genes shown on the left. Protein levels were normalized to the maximal level across all FACS gates. **b**, Periportal bias in expression of mRNA and proteins, calculated as the difference between the two periportal gates and the two pericentral gates, normalized by the mean expression across all gates. Light grey – all matched mRNA and proteins. Dark grey – mRNA and proteins with minimal expression fraction higher than 10^−5^ in any of the gates for both mRNA and protein. Spearman’s r is indicated for each dataset. Dashed line marks a slope of 1. **c**, Expression profiles of mRNA (grey) and their respective proteins (red). Mean of five mice is plotted. Error bars represent SEM.

The combined measurements of both mRNA and proteins from the same spatially-sorted gates enabled a controlled comparison of protein and mRNA zonation patterns (Fig. b-c). The periportal biases (the difference between the expression in the periportal and pericentral gates divided by the mean expression) were significantly correlated between mRNA and proteins, indicating similar mRNA and protein zonation profiles for most genes (Spearman’s r = 0.39, p-val = 1.45×10^−110^, for protein and mRNA with minimal expression level higher than 10^−5^ r=0.45, p-val = 1.71×10^−79^ Fig. b-c). Notably, some genes exhibited discordant zonation of mRNAs and proteins. These included genes that were zonated at the protein but not mRNA level, such as Rbp4, Idh3b, Mrpl43 and genes that were zonated at the mRNA but not at the protein level, such as A1cf, Clmn and Lsr (Fig. 5c). The discordant genes also included Hnf4a, a key hepatocyte transcription factor^4,17^. The mRNA levels of Hnf4a were not zonated, whereas the protein content was higher in the periportal gates. This periportal protein bias is in line with previously reported involvement of Hnf4a in periportal repression of Wnt regulated pericentral genes^17–19^ and induction of periportally expressed targets^20^. Thus, our analysis indicates that the majority of proteins and mRNAs are similarly zonated, and highlight genes with potential post-transcriptional regulation.

### Zonation of the hepatocyte microRNA content

We next asked whether spatial sorting could be used to explore the regulatory mechanisms that shape hepatocyte zonation. MicroRNAs (miRNAs/miRs) are short RNA oligonucleotides, roughly 22 bp long, that target specific mRNAs through Watson-Crick base-pairing, leading to increased degradation or decreased translation of target transcripts^21^. Regulation by miRs seems to be important in liver development, metabolism and homeostasis ^22,23^. Notably, miR regulation may impact liver zonation, as mice lacking the miR central processing element Dicer in hepatocytes exhibit profound changes in zonation patterns^24^. We reasoned that combined global measurements of the zonation profiles of miRs and mRNA could identify potential miR-target regulatory interactions through the detection of miR-target pairs with anti-correlated expression profiles^25^.

To this end, we performed microRNA microarray measurements on spatially sorted hepatocytes from three mice. We detected 302 miRs that were expressed in hepatocytes in all three mice. We further focused on 137 miRs that were classified as “high confidence” in miRBase^26^. 45% (61/137) of these high-confidence hepatocyte-expressed miRs were significantly zonated (FDR ≤ 0.2, Kruskal-Wallis test with Benjamini-Hochberg correction, Fig. 6a). Most zonation profiles (48/61) were mildly pericentral with 4 ≤ COM ≤ 4.5 (COM = center of mass, Methods), while seven others showed strong periportal zonation (COM ≥ 6). We measured the expression of six of the miRs predicted to be zonated using qRT-PCR, obtaining excellent correspondence with the microarray measurements (mean r_Pearson_ = 0.83±0.23, Fisher’s method *p*< 10^−37^, Fig. 6b, Methods).

**Fig 6.**
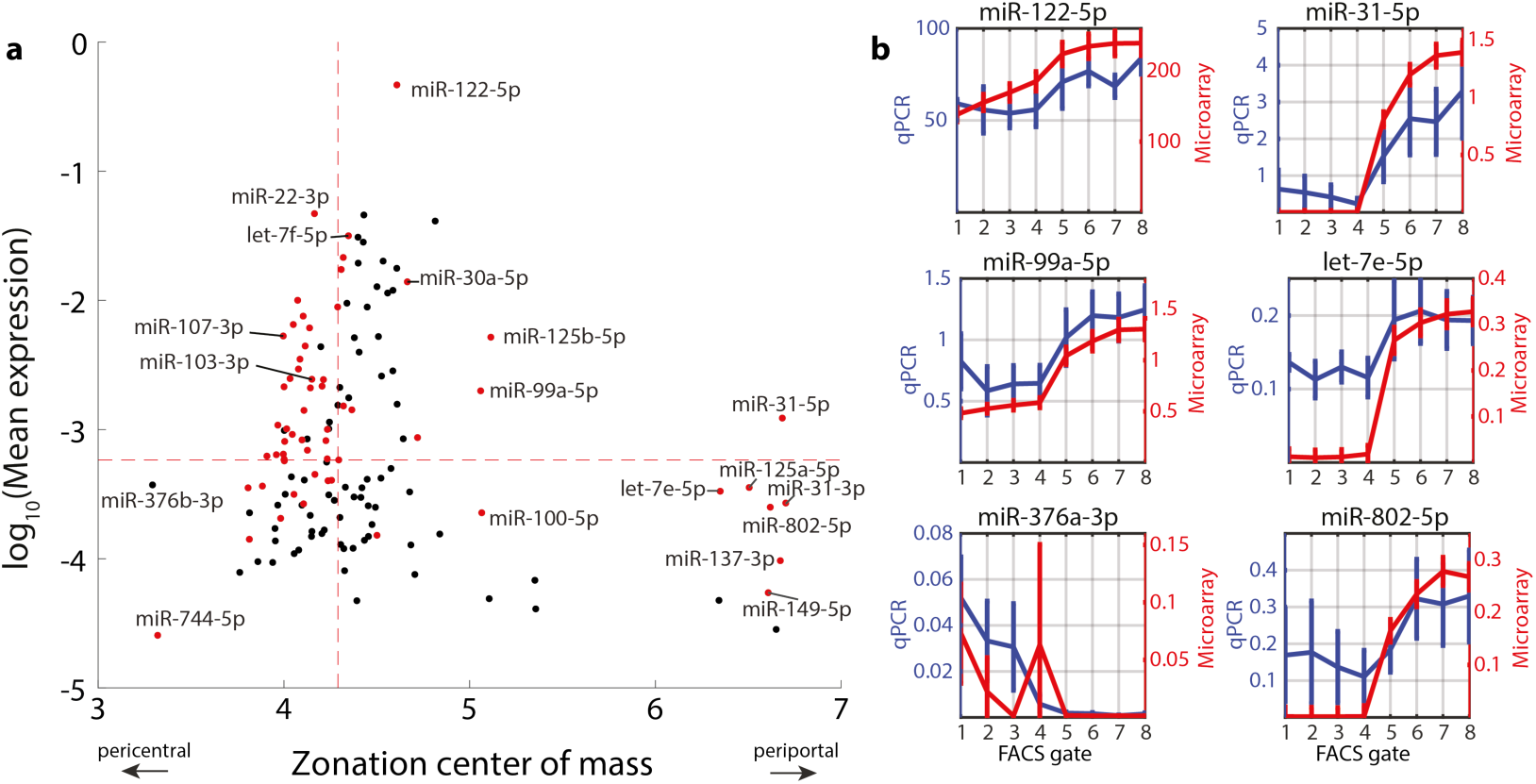
Zonated expression of hepatocyte miRs. **a**, Mean expression vs. zonation profile center of mass for all detected high-confidence miRs. Selected miRs are labelled. Dashed red lines denote the median of each quantity. Red dots are miRs that are significantly zonated (FDR ≤ 0.2). **b**, Validations of hepatocyte miR zonation profiles using qRT-PCR. Profiles for both qRT-PCR and microarrays are normalized by expression levels of miR-103-3p (r_Pearson_ = 0.83±0.23, Methods). Error bars indicate SEM. Discrepancies between qRT-PCR and microarray profiles for let-7e-5p, miR-376a and miR-802-5p may be due to limited sensitivity of the microarray at low expression levels.

The zonated miRs included miRs previously described to play a role in liver development, metabolism and regeneration. MiR-122-5p, the most abundant miR in our measurements, in agreement with previous studies^27^, comprised 46.5±3.5% of the total miR content in hepatocytes. We found that miR-122-5p was periportally zonated, with a 1.15-fold higher expression in the periportal gates compared to the pericentral gates (p-value < 0.01, Kruskall-Wallis test). miR-122-5p was significantly anti-correlated with its targets compared to randomized genes (Methods), indicating a potential regulatory role in shaping their zonation. Prominent pericentral miR-122-5p targets (genes that were repressed in their expression in the periportal layers in which miR-122 was more abundant) included the canonical miR-122-5p target gene Cs, encoding citrate synthase, as well as Klf6 and Slc35a4^28^ (Supplementary Fig. 4). MiR-30a-5p exhibited periportal zonation (periportal to pericentral ratio of 1.19, p=0.007, Fig. 6a, Supplementary Table 4). Mtdh, a known target of miR-30a-5p, previously shown to change in expression in liver tumors^29^, was pericentral, inversely zonated to its miR regulator (r_Spearman_ = −0.81, p = 0.022, Supplementary Table 5). Additional zonated miRs included the pericentral miR-103-3p and miR-107-3p and the periportal miR-802-5p, which have been previously shown to modulate hepatic glucose sensitivity^30,31^ (Fig. 6a). In summary, our measurements revealed profound zonation of key hepatic miRs.

### Computational approach for detection of putative miR-regulated hepatocyte target genes using zonation profiles

Spatially-stratified measurements of miRs and mRNAs could be use identify potential miR regulation at the mRNA degradation level. Such regulation would be manifested in inverse correlations between the zonation profiles of a target mRNA and its regulating miR(s). To identify such interactions, we constructed a miR-mRNA regulatory network based on predictions from TargetScan^32^ (Supplementary Table 5). We included all hepatocyte-expressed genes and interactions with high confidence (Methods). The resulting network included 33,672 interactions between 131 miRs and 6,650 genes. For each gene, we constructed the cumulative regulating miR profile, by summing up the zonation profiles of all miRs with a predicted regulatory interaction for the considered target gene (Supplementary Table 6). We computed the Spearman correlation between the gene’s mRNA zonation profile and the cumulative miR zonation profile and compared it to randomized degree-preserving networks (Fig. 7a, Methods).

**Fig 7.**
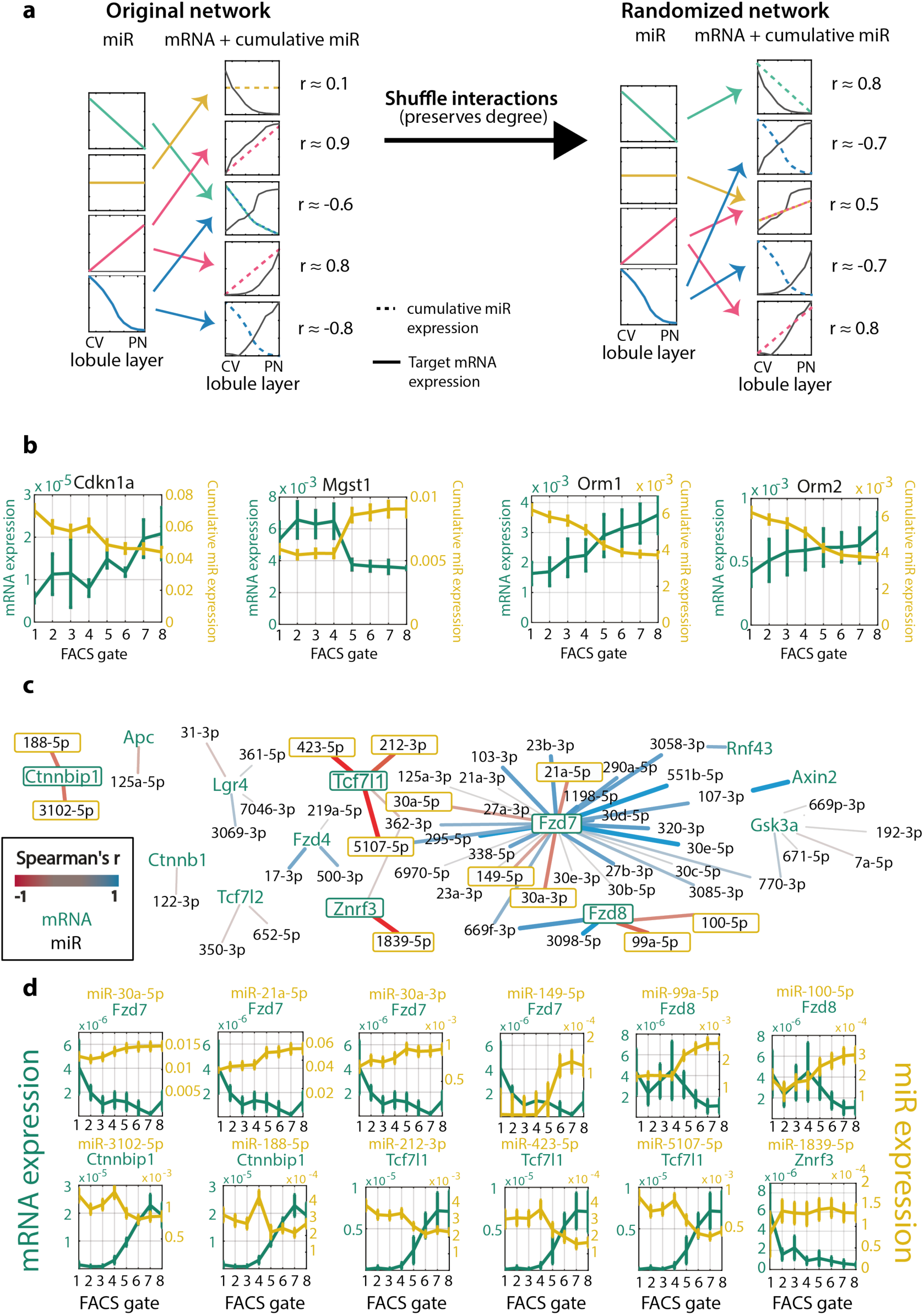
Network analysis of miR-target interactions. **a**, Schematic illustration of the algorithm for inferring significant interactions between miRs and target genes. **b**, Zonation profiles of selected genes and their significantly anti-correlated cumulative miR profiles. **c**, Regulatory network of hepatocyte-expressed Wnt pathway components and their expressed regulating miRs. Edges colored by the correlation between the miR and target. Edge weight is proportional to the absolute correlation value. **d**, Selected pairs of miRs and regulated Wnt signaling components (outlined in green in Fig. 7c). The transcripts of Ctnnbip1, Fzd8, Tcf7l1 and Znrf3 are anti-correlated with most of their regulating miRs, suggesting that miRs have a relatively more important role in regulating these genes’ expression in comparison to other genes.

Our analysis identified 45 genes that were significantly more anti-correlated with their regulating miRs compared to random (FDR ≤ 0.2, Supplementary Table 6). Pericentral target genes included Mgst1, which encodes the enzyme microsomal glutathione S-transferase that conjugates glutathione to hydrophobic electrophiles. Periportal target genes included cyclin-dependent kinase inhibitor 1 (also known as p21), encoded by Cdkn1a. Cdkn1a showed anti-correlation with 10 of its 11 regulating miRs (median r_Spearman_ = −0.79, Supplementary Fig. 5), including miR-20a-5p, miR-20b-5p, miR-22-3p miR-93-5p and miR-106b-5p that were experimentally validated as Cdkn1a regulators^33,34^. Alpha-1-acid glycoprotein 1 (AGP1), and Alpha-1-acid glycoprotein 2 (AGP2), encoded by the genes Orm1 and Orm2 respectively, are secreted plasma carrier proteins that were periportal target genes, significantly anti-correlated with all of their regulating miRs miR-20a-5p, miR-20b-5p, miR-93-5p and miR-106b-5p (mean r_Spearman_ = −0.994±0.011, Fisher’s method p < 10^−14^, Fig. 7b, Supplementary Fig. 5). These miRs, which also regulate Cdkn1a, are considered part of the same “miR family” due to their high seed sequence similarity^33^ and exhibit similar zonation patterns (Supplementary Fig. 5), suggesting modular expression and activity of this miR family.

### Regulation of Wnt signaling components by miRNA

Wnt is a major factor that shapes hepatocyte zonation^35–40^. Wnt and Rspondin morphogens are secreted by pericentral liver endothelial cells^2,11,41–43^, resulting in higher pericentral expression of Wnt-activated genes and lower pericentral expression of Wnt-inhibited genes^5,35^. Notably, hepatocyte-specific Dicer knock-out mice have perturbed zonation of Wnt-regulated genes, such as Glul and Arg1^24^. This suggests that miRs could differentially modulate hepatocyte Wnt signaling in different lobule zones. To explore this hypothesis we analyzed the miR-target sub-network that includes genes associated with Wnt signal processing (Methods, Fig. 7c-d). This analysis uncovered several key components of the Wnt network that exhibit zonation in hepatocytes and that have spatially-anticorrelated zonated miRs. The Wnt receptors Fzd7 and Fzd8 were more highly expressed in pericentral hepatocytes, whereas their regulating miRs miR149-5p, miR-30a-5p, miR-30a-3p, miR-21a-5p, miR-99a-5p and miR-100-5p were more abundant in periportal hepatocytes. In contrast, inhibitory components of Wnt signaling such as Ctnnbip1 and Tcf7l1 were periportally zonated. Tcf7l1, also known as Tcf3, is a transcriptional repressor of Wnt-activated genes that is inactivated by binding of β-catenin^44^. This periportally zonated gene is anti-correlated with its regulators miR-212-3p, miR-423-5p and miR-5107-5p (Fig. 7d). Ctnnbip1, encoding β- catenin interacting protein, prevents the binding of β-catenin to TCF7L1 and thus its removal and activation of Wnt target genes^45^. The miR regulators of this periportal gene, miR-188-5p and miR-3102-5p, were pericentrally-zonated (Fig. 7d). Our analysis thus highlights zonated components of hepatocyte Wnt signaling and their potential regulation by miRs.

## Discussion

The liver exhibits profound division of labor among hepatocytes that reside at different zones. Understanding and modeling liver function thus requires characterizing the hepatocyte functions at each lobule coordinate. In this study, we present spatial sorting, a generic approach to isolate large amounts of hepatocytes with high spatial resolution for a broad range of downstream measurement modalities. The approach utilizes zonated surface markers that can be identified by spatially resolved transcriptomic atlases. We demonstrated applications of this approach for resolving the zonation of hepatocyte proteins and miRs. The approach can be readily applied to other structured organs and cells types exhibiting zonation, including liver endothelial cells^11^, intestinal enterocytes^46^ and kidney cells^47,48^. The usage of endogenous surface markers renders spatial sorting particularly useful for studying zonation in humans as well.

Our proteome analysis revealed some notable discordance between the average hepatocyte levels of proteins and mRNAs, mostly for genes encoding secreted proteins (Fig. 4). In contrast, we found that the protein zonation profiles highly overlap those of the mRNAs. These results argue for a predominance of spatial regulation of hepatocyte protein levels via transcription or mRNA stability, rather than through translational control or protein stability. Hnf4a, a key hepatic transcription factor, was among the small group of genes for which protein and mRNA zonation profiles were discordant. Hnf4a mRNA was expressed in a non-zonated manner, whereas its protein levels were periportally zonated. This fits with previous reports of periportal expression of Hnf4a hepatocyte target genes^4,18–20^. Notably, Hnf4a is a transcriptional activator of miR-122^49^, the most abundant liver-expressed miR, which we also found to be periportally zonated. Due to sensitivity limitations of mass-spectrometry proteomics our proteomic measurements did not include low-abundance genes, including other key liver transcription factors, which may exhibit higher levels of post-transcriptional regulation.

Recent works have begun to develop in-silico multi-scale models for predicting the liver’s response to stimulations by metabolites and xenobiotics^50–52^. These models consider multiple units representing hepatocytes at different zones that exchange materials and process them through individualized metabolic networks, thus modeling the polarized blood perfusion throughout the lobule. Future incorporation of the zonated levels of enzymes into such models could increase their precision and better capture in-vivo fluxes. Our proteomic map provides such detailed zonation of key enzymes (Supplementary Table 3).

Our work provided a global spatial atlas of miR zonation, identifying key hepatocyte zonated genes such as miR-122-5p and miR-30a-5p. We used the combined miR and target mRNA levels to identify potential regulatory interactions that could entail zonated mRNA degradation. This forms an important resource for future functional validations. MiRs have been shown to be highly dynamic along the course of several diseases such as fibrosis, viral infection and liver cancer^23,53,54^. Spatial sorting could be used to measure the zonation of these miRs along the courses of these diseases. Moreover, plasma measurements of zonated miRs^55^ could potentially be used as biomarkers to identify zonated liver damage.

Wnt is a key regulator of hepatocyte zonation^2,35,39^. The pericentral secretion of Wnt morphogens by endothelial cells generates a zonated external morphogen field, that could explain a significant fraction of the zonated hepatocyte genes. Our study revealed that in addition to the Wnt signal, the hepatocyte Wnt sensing and processing machinery also seems to be zonated. We measured pericentral expression of the key Wnt receptors Fzd7 and Fzd8 and periportal zonation of the Wnt inhibitors Tcf7l1 and Ctnnbip1. This joins previous reports of periportal zonation of APC, a key Wnt regulator^35^. Our study further identified miRs that regulate these zonated Wnt components. Thus miRs seem to be upstream of Wnt signaling. These results could explain the effects of hepatocyte-specific Dicer KO and beta-catenin KO. While Dicer KO resulted in perturbed zonation of Wnt targets, beta-catenin KO did not substantially alter miR levels^24^.

Our approach enables attaining up to a few hundreds of thousands of hepatocytes per sorted population. While this amount is compatible with a broad range of assays, it is insufficient for assays that require massively larger amounts of material, such as RNA methylations^57^ and metabolic profiling^58^. Moreover, since the approach is FACS-based, measuring metabolites, which are labile, would be compromised by the substantial incubation periods involved in the protocol^59^. Nevertheless, it will be interesting to apply spatial sorting to explore additional zonated hepatocyte features, including chromatin modifications, DNA methylations, three-dimensional chromosomal conformations, DNA mutation spectra and chromosomal aberration. Such measurements will resolve hepatocyte cell identity, regulatory mechanisms and susceptibility to damage in each zone.

## Acknowledgements

We thank Efrat Hagai and the Flow Cytometry Unit (Weizmann Institute of Science) for the FACS technical support; Dr. Tamar Ziv and the Smoler Proteomics Center (Technion) for performing the LC-MS/MS and analyzing the results; Dr. David Pilzer and the Genomic Technologies Unit (Weizmann Institute of Science) for performing the miR microarray measurements. We thank all members of the lab for valuable comments. S.I. is supported by the Henry Chanoch Krenter Institute for Biomedical Imaging and Genomics, The Leir Charitable Foundations, Richard Jakubskind Laboratory of Systems Biology, Cymerman-Jakubskind Prize, The Lord Sieff of Brimpton Memorial Fund, the I-CORE program of the Planning and Budgeting Committee and the Israel Science Foundation (grants 1902/ 12 and 1796/12), the Israel Science Foundation grant No. 1486/16, the EMBO Young Investigator Program and the European Research Council under the European Union’s Seventh Framework Programme (FP7/2007-2013)/ERC grant agreement number 335122, the Bert L. and N. Kuggie Vallee Foundation and the Howard Hughes Medical Institute (HHMI) international research scholar award.

## Author Contributions

K.B.H. and S.I. conceived the study, S.B.M. and S.I designed experiments, S.B.M. prepared all the samples, S.B.M and Y.S. analyzed the data, K.B.H contributed to establishing the method, A.E.M contributed to data analysis, S.I. supervised the study, S.B.M, Y.S. and S.I. wrote the manuscript. All authors reviewed the manuscript and provided input.

## Conflicts of Interest

The authors declare no competing interest.

## Methods

### Animal experiments

Mouse experiments were approved by the Institutional Animal Care and Use Committee of the Weizmann Institute of Science and performed in accordance with institutional guidelines. Sorting experiments were conducted on five three months old C57BL/6 male mice, obtained from Harlan laboratories. Mice were fed ad libitum and were kept in a reverse light-dark cycle. Mice were anaesthetized with Ketamine (100 mg kg−1) and Xylazine (10 mg kg−1) dissolved in 1xPBS and injected intraperitoneally 6h-9h after lights off (ZT 18-21).

For imaging experiments, livers of 3m old wt male mice were harvested and fixed in cold PFA for 3h at 4°C followed by overnight fixation in cold PFA + 30% sucrose at 4°C while revolving. Livers were embedded in OCT (Tissue-Tek) the next morning. Blocks were kept at −80°C.

### Immuno-Fluorescence

OCT embedded mouse liver blocks were sectioned into 7um thick slices. Slices were fixed with cold Methanol for 20min. After three 5min washes with PBST (1X PBS, 1% BSA + 0.1% Tween), sections were permeabilized by 10min incubation at room temperature with PBSTX solution (1X PBS, 0.25% Triton 100X and 1% BSA). Slices were then washed again as before and were incubated for 1h at room temperature with blocking solution (1x PBS, 0.1% Tween and 5% Goat/Normal Horse Serum). Slices were next incubated with the antibody solution (blocking solution with 1:50 antibody in a total reaction volume of 150ul) at 4°C overnight. Antibodies used were Alexa Fluor 647 rat anti-mouse CD73 (BD, cat: 561543) and Alexa Fluor 555 mouse anti-Ecadherin (BD, cat: 560064). On the next day, slices were washed with PBST 3 times. Nuclei were stained with DAPI (1:100 in PBS, 10mis). Imaging of liver porto-central axis was performed on a Nikon-Ti-E inverted fluorescence microscope with a 100× oil-immersion objective and a Photometrics Pixis 1024 CCD camera using MetaMorph software using the scan stage option.

Z-projected images of lobule scans (8 scans, 3 mice) were analyzed. Membrane segments of hepatocytes were measured for the intensity of Alexa Fluor 555 (E-cadherin) and Alexa Fluor 647 (CD73). Background, set as the paired cytoplasmic intensity for each membrane signal was subtracted. Segments were then binned into eight groups representing eight lobule layers (1 = pericentral, 8 = periportal), according to their radial distance from the central vein. Median intensity of the segments from each lobule layer was calculated and averaged over the different lobules (Fig. 2b-c). Values were scaled from 0 to 1 and the plot was smoothened with a sliding window of 3.

### Liver Perfusions and hepatocytes dissociation

Once anaesthetized, mice livers were perfused as previously described^60^, with a few adjustments. A 27G syringe, connected to the perfusion line and pump, was inserted into the vena cava. 25ml of pre-warmed to 37°C EGTA buffer followed by 25ml of pre-warmed to 37°C EBS buffer with 2.3U of Liberase Blendzyme 3 recombinant collagenase (Roche Diagnostics) were cannulated into the vena cava. Shortly after the beginning of the perfusion, the portal vein was cut to allow drainage of the blood.

After perfusion, livers were explanted into a Petri dish with 25ml of pre-warmed EBS and gently minced using forceps. Dissociated liver cells were collected and filtered through a 100um cell strainer. Cells were spun down at 30rcf for 3 min at 4°C to get hepatocytes enriched pellet. Pellet was resuspended in 25ul cold EBS.

### Cells Staining

To discard dead hepatocytes, 22.5ml Percoll (Sigma) mixed with 2.5ml 10x PBS was added to the cells. Cells were centrifuged at 600rpm for 10 minutes. Supernatant containing the dead cells was aspirated and cells were resuspended in pre-warmed Hoechst buffer (DMEM + 10% FBS + 10mM Hepes). After counting, concentration was adjusted to 2×10^6^ cells in 1ml. To determine ploidy of hepatocytes, DNA was stained with Hoeschst (15ug/ml). Resperine (5uM) was also supplemented to the cells to prevent Hoechst expulsion from the cells. Cells were incubated 30min at 37°C. Hepatocytes were centrifuged for 5min in 1000rpm at 4°C and supernatant was discarded. Next, cells were stained with Alexa fluor 488 Zombie green (BioLegend) to later enable the detection of viable cells by FACS. Cells were resuspended in cold PBS in a concentration of 10°cells in 100ul. Zombie-green was added in a dilution of 1:500. Cells were kept in a rotator in the dark at room temperature for 15min. After spinning down (1000rpm, 5min, 4°C), cells were resuspended in FACS buffer (2mM EDTA pH 8 and 0.5% BSA in 1xPBS), in a concentration of 10^6^ cells in 100ul. Cells were stained with PE-anti-E-cadherin (BioLegend, cat: 147304), APC-anti-CD73 (BioLegend, cat: 127210), PE-Cy7-anti-CD31 (BioLegend, cat: 102418) and APC-Cy7-anti-CD45 (BioLegend, cat: 103116), in a dilution of 1:300. FcX blocking solution (BioLegend) was added in a dilution of 1:50.

### Flow Cytometry and sorting

Cells were sorted by SORP-FACSAriaII sorter (BD) using a 130 μm nozzle and 1.5 natural density (ND) filter. Lasers compensation was corrected manually. In order to collect eight populations, each enriched with spatially-stratified hepatocytes with equal viability and ploidy levels, events were screened through the following five nested gates (Fig. 3a-b): (1) hepatocytes gate from all events – set by plotting FSC-A against SSC-A and excluding large clusters and small debris; (2) singlets FSC – set by excluding the margins of FSC-A and FSC-W plot; (3) singlets SSC – excluding upper margins of SSC-W when plotted against SSC-A; (4) live cells gates according to the Zombie-488 negative cells, comparable to unstained cells; (5) hepatocytes only, by depleting CD31 and CD45, NPCs markers, and (6) tetraploid hepatocytes, inferred by Hoechst histogram (Fig. 3a-b, Supplementary Fig. 1b-c). Hepatocyte size and overall protein content scale with ploidy^61^, thus creating spurious correlations between the zonated surface markers (Supplementary Fig. 1). Sorting without ploidy stratification would result in inclusion of hepatocytes from different lobule layers, reducing spatial accuracy (Supplementary Fig. 1).

We then plotted PE-intensity for E-cadherin staining and APC-intensity for CD73 staining. The positively stained cells were determined by measuring the intensities for unstained cells. The highest intensity for unstained cells was the threshold for the positively stained cells. Each population, CD73 positive and E-cadherin positive, was further gated to four equal subpopulations, representing graded intensities of the marker. Thus, subpopulations 1, 2, 3 and 4 had equal amount of events, 1 had the highest APC-CD73 intensity while 2, 3, 4 have gradually decreasing intensities of APC. Likewise, subpopulations 5, 6, 7 and 8 were equally distributed, 8 has the highest PE-E-cadherin intensity while 7,6,5 have gradually decreasing PE intensities. Populations 4 and 5 contained cells from below positive intensity threshold, to accurately resemble mid lobule hepatocytes, in which both CD73 and E-cadherin abundances are very low (Fig. 2). All gates were set for each of the five experiments independently, with a large overlap.

10,000 hepatocytes from each gate were sorted into Dynabeads mRNA DIRECT Micro Kit lysis buffer (Invitrogen) for RNA sequencing. After sorting, cells were spun down and frozen at −80°C until processing. 100,000 hepatocytes from each population were collected into FACS buffer, and resuspended twice with PBS to wash away serum proteins. Pellets were flash frozen and sent to Mass-spectrometry proteomics measurements (The Smoler Protein Research Center, Technion, Israel). Additional 50,000 cells were collected for microRNA microarray. Total RNA was isolated using Direct-zol RNA microprep kit (Zymo Research).

### RNA sequencing

10,000 hepatocytes from each sorted population were collected for library preparation. Cells were sorted into Lysis buffer supplied in Dynabeads mRNA Purification Kit (Invitrogen, cat: 61006). RNA was extracted by the kit according to the provided protocol. 2ul of the extracted mRNA from each sample were used for libraries. Library preparation was done with mcSCRBseq protocol^62^. The cDNA was pre-amplified with 10-15 cycles, depending on cDNA concentration indicated by qPCR quality control. 2ng of the amplified cDNA was converted into sequencing library with the Nextera XT DNA Library kit (Illumina, FC-131-1024), according to supplied protocol. Quality control of the resulting libraries was performed with an Agilent High Sensitivity D1000 ScreenTape System (Agilent, 5067-5584). Libraries were loaded with a concentration of 2.2pM on 75 cycle high output flow cells (Illumina, FC-404-2005) and sequenced on a NextSeq 500 (Illumina) with the following cycle distribution: 8bp index1, 16 bp read1, 66 bp read2 (no index2 needed). Total 40 libraries, eight sorted populations for five different mice were sequenced.

### Sequencing analysis pipeline

Illumina output files were demultiplexed with bcl2fastq 2.17 and the resulting fastq files of mRNA sequencing experiments were analyzed with the zUMIs pipeline^63^. Reads were aligned using STAR to a transcriptome index of the GRCm38 release 84 (Ensembl) and exonic UMI counts per million (CPM) were calculated with pipeline default settings and TMM normalization^64^ implemented in EdgeR^65^. 14,027 transcripts were identified in the experiment over the 40 libraries. Data were further normalized by dividing each sample by its sum of CPM. Two out of the 40 libraries failed to reach over 200K reads and were discarded (m2_2_cpm and m3_5_cpm samples in Supplementary Table 1).

### Mass spectrometry

Fourty samples (five mice, eight populations each) were digested by trypsin and analyzed by LC-MS/MS on Q Exactive plus (Thermo). The data was analyzed with MaxQuant 1.5.2.8^66^ against the mouse Uniprot database. Data were quantified using the same software. We retained proteins with FDR <0.01 in at least 2 samples in one of the eight groups, identified by at least 2 peptides across all samples. 3,210 proteins were identified. For each sample, LFQ intensities for each peptide were normalized by the sum of all intensities – yielding expression fraction out of the total protein detected.

### MiRNA Microarrays

Total RNA (100ng), isolated from bulk populations of 50,000 spatially-sorted hepatocytes (*n* = 3) per FACS gate, was labeled with Cy3 during transformation into cDNA using an RNA Agilent miRNA Labeling Kit (Agilent, UK) and Spike Kit (Agilent, UK). cDNAs were hybridized to Mouse miRNA Microarray, Release 21.0, 8×60K (v21) microarray slides (Agilent, UK) according to Agilent microRNA Hybridization Kit protocol (Agilent, UK) and scanned using Agilent G2505B array scanner. Data were extracted using the Feature Extraction software (Agilent, UK) with default parameters.

### qRT-PCR

Total RNA, isolated from bulk populations of 50,000 spatially-sorted hepatocytes (*n* = 3) per FACS gate, was diluted to 5 ng/µL, and cDNA was reverse-transcribed using the miRCURY LNA RT Kit (Qiagen, cat. no. 339340) according to the manufacturer’s instructions on an Applied BioSystems ProFlex PCR System. Plates were prepared using the miRCURY SYBR Green PCR Kit (Qiagen, cat. no. 339346) with custom miRCURY LNA PCR primers (Qiagen cat. no. 339306, Supplementary Table 7). Each 10 µL reaction volume contained 5 µL 2x miRCURY SYBR Green Master Mix, 0.5 µL ROX reference dye, 1 µL PCR primer mix, 0.5 µL RNAse-free water and 3 µL of cDNA sample diluted 1:60. qPCR reactions and measurements were performed on a StepOne Real-Time PCR System (Thermo Fisher, cat. no. 4376357) according to the manufacturer’s instructions. Relative expression levels were calculated as

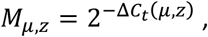

where *M* _*µ, z*_ is the relative expression of miR *µ* in FACS gate *z*, δ*C_t_ = C* _*µ, z*_- *C _n,z_ C* _*µ, z*_ is the *C_t_* (threshold cycle) of miR *µ* in FACS gate *z* and *C_n,z_*is the *C_t_* value of miR-103-3p in FACS gate *z* (isolated from the same mouse).

### Center of mass calculation

The center of mass (COM) of an expression profile *x* (spread over *z =* 1: 8 FACS gates) was calculated as:

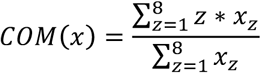

This formula yields a number *COM* ∈ [1,8] that indicates around which gate most of the expression is distributed.

### Comparing bulk mRNA with published scRNAseq

Spatial sorting produces sub-populations of hepatocytes that are enriched for specific lobule layers, however each FACS gate includes several lobule layers. To compare the bulk mRNA measurements of the FACS-gated sub-populations to the zonation measurements previously reconstructed using spatially-resolved single cell transcriptomics^5^ we thus computationally estimated the center of mass, namely the weighted average of all single cells represented by each gate. To this end, we implemented Cibersort (https://cibersort.stanford.edu/^67^, to estimate the relative abundances of each lobule layer in the sorted subpopulations. We extracted a gene signature list for each layer from the scRNA seq data^5^. A total of 17 genes with mean expression greater than 5×10^−5^, zonation FDR smaller than 0.01 and dynamic range of at least 10-fold between the mean of the two periportal layers and the mean of the two pericental layers was used for the analysis. The means of five mice of the zonation profiles of these genes across the eight sorted gates were used as the mixed-populations data set. The relative abundances of each of the nine layers in each of the eight sorted population was calculated (with ‘Disable quantile normalization’ option checked). Fig. 3c presents the mean expression in each FACS gate over five mice (blue) and mean expression in scRNA seq data^5^, weighted by the relative abundances of each layer in each FACS gate (yellow).

### Comparing proteins and RNA

Out of the 3,210 proteins (Supplementary Table 1) detected in Mass-spectrometry and 14,027 mRNAs (Supplementary Table 2) detected in RNA-sequencing, 3,051 were found in both datasets (Supplementary Table 3). The median expression fraction of five mice was calculated for each gate in each measurement. We scatter-plotted the averages over all gates of the eight mRNA medians and eight protein medians for every gene and found a Spearman correlation r= 0.50 (0.48-0.50 per each gate independently). In order to better characterize mRNA and protein ratios in different KEGG pathways^68^, we plotted the regression line of protein by mRNA. The residual of the proteins from the regression line was calculated and grouped according to KEGG pathways (Fig. 4b, Supplementary Fig. 2).

### Computing zonation

For each of the 3,051 common proteins and mRNAs, Kruskal-Wallis test was performed to check for variability between different sorted gates. To correct for multiple hypotheses, we performed the Benjamini–Hochberg procedure to obtain the FDR for each hypothesis. We classified proteins as zonated if they had FDR < 0.05. 1,672 were significantly zonated. To produce the protein zonation heatmap (Fig. 5a), we first removed all proteins which have a median of LFQ = 2^18^ in any of the eight gates (479 proteins). Next, we normalized all protein profiles to their maximum across all FACS gates and sorted them by their center of mass (Fig. 5a).

### Statistical analysis of miR data

*Microarray data:* only miRs that were annotated as “high confidence” in miRBase^26^ v22 (downloaded 30/10/18) were kept for analysis. The raw signal for each miR in each FACS gate and in each array was normalized by the total signal per gate per array. Only miRs present in all three biological replicates were retained for further analysis, and their initial normalized signal was averaged over all arrays. Finally, the averaged signal was divided again by the total signal in each gate (this operation amounted to dividing by a number very close to 1, since only miRs with very low expression were not present in only some of the replicates). MiR zonation was inferred using the Kruskall-Wallis test (for each miR, comparison of 8 gates, with each having 3 replicates), and applying a Benjamini-Hochberg correction on the p-values obtained from the KW test. MiRs with FDR ≤ 0.2 were classified as zonated.

#### Differential zonation of miR-122-5p targets

Targets of miR-122-5p were taken from ref.^28^. 146 of the targets listed were expressed in our liver zonated transcriptome data. The mean COM was calculated for these targets and for 1,000 random samplings (with replacement) of 146 liver-expressed genes (genes with maximal expression over all gates that was at least 4×10^−6^ of transcriptome). We performed a Wilcoxon rank-sum test for the COMs of the target genes vs. COMs of randomly sampled genes, yielding p = 0.018.

#### MiR-target network construction, randomized networks and genes more anti-correlated with their cumulative miR profiles than random

all miR-target interaction predictions were downloaded from TargetScanMouse v7.2^32^ (data released 8/2018, downloaded 24/10/18), including conserved and non-conserved sites. Predicted edges were filtered for only liver-expressed miRs, annotated “high confidence” in miRBase v22, weighted context++ score percentile ≥ 95 and genes expressed with maximum fraction (over gates) of total transcriptome ≥ 4×10^−6^). The resulting network included 33,672 interactions between 131 miRs and 6,650 genes. For each gene *g* we constructed the cumulative expression profile *M*_*g*_(*z*) of all miRs *µ*_*i*_ (*z*), *i* ∈ {1,*2, …*, *N*_*g*_, *z =* {1, *…* 8} predicted to target it:

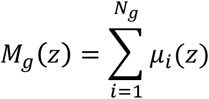

and calculated the Spearman correlation between the expression profile *X*_*g*_(*z*) of each gene *g* and *M*_*g*_(*z*). We then created 1,000 networks with randomized edge assignment using mfinder^69^ with the command *mfinder -r 1000 -ornet*. We took into account miRs that regulate the same target genes at multiple sites, and for the purpose of network randomization, these were considered as separate edges by creating “virtual” miR nodes that were collapsed back to the original miR after randomization. For each randomized network, we calculated again the cumulative miR profile for each of the 6,650 genes and the corresponding Spearman correlation. We then calculated for each gene the fraction of randomized networks in which the anti-correlation of the gene and the original predicted cumulative miR expression profiles is smaller than the anti-correlation of the gene with the cumulative profiles generated with the randomized networks, generating an empirical p-value *p*. We manually corrected genes with *p=* 0 to *p*→ *p*′*=* 1/*N*, with *N* = 1,000 the number of networks generated. FDR values using the Benjamini-Hochberg procedure were calculated for all p-values and genes with FDR ≤ 0.2 were deemed “significant” (Supplementary Table 6).

#### Regulation of Wnt pathway components

we examined all edges in our miR-target network that included genes which are involved in Wnt signaling transduction^70^, and that were anti-correlated with individual miRs regulating them with a Spearman coefficient of −0.5 or less. The genes were Apc, Axin2, Ctnnb1, Ctnnbip1, Dvl1/2, Fzd1-10, Gsk3, Lgr4/5/6, Lrp5/6, Rnf43, Tcf7, Tcf7l1/2 and Znrf3.

#### Network visualization

the miR-Wnt pathway component network was visualized using Cytoscape v3.7^71^. All detected Wnt pathway components with miRs predicted to regulate them (see “miR-target network construction”), and the Spearman correlations between them, were used as input.

## Supplementary Figures

**Supplementary Fig. 1.**
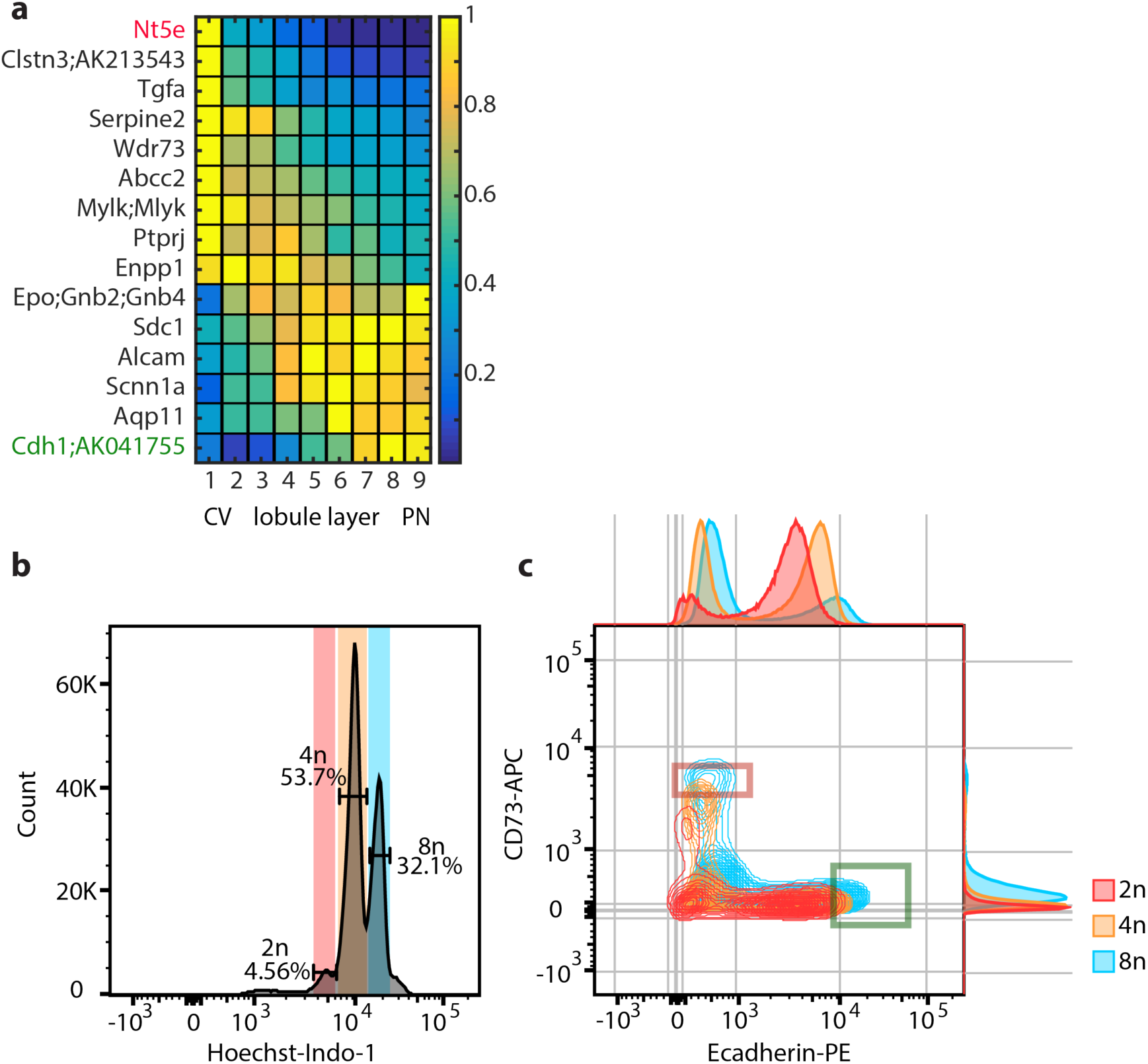
Surface markers candidates for spatial sorting and ploidy gating in FACS. **a**, Surface markers zonation profiles. Data from ^5^. Expression is normalized to the maximal level for each gene. Genes are sorted by expression center of mass. **b-c**, Gating by ploidy levels. **b**, Histogram of cells according to Hoechst stains, proportional to DNA content, each peak represents a different ploidy class with different proportions in the hepatocytes population. Red – diploid cells, orange – tetraploid cells, blue – octoploid cells. **c**, distribution of hepatocytes according to E-cadherin and CD73 intensities, stratifies by ploidy class. Red and green rectangles are 1 and 8 gates selected for the 4n hepatocytes.

**Supplementary Fig. 2.**
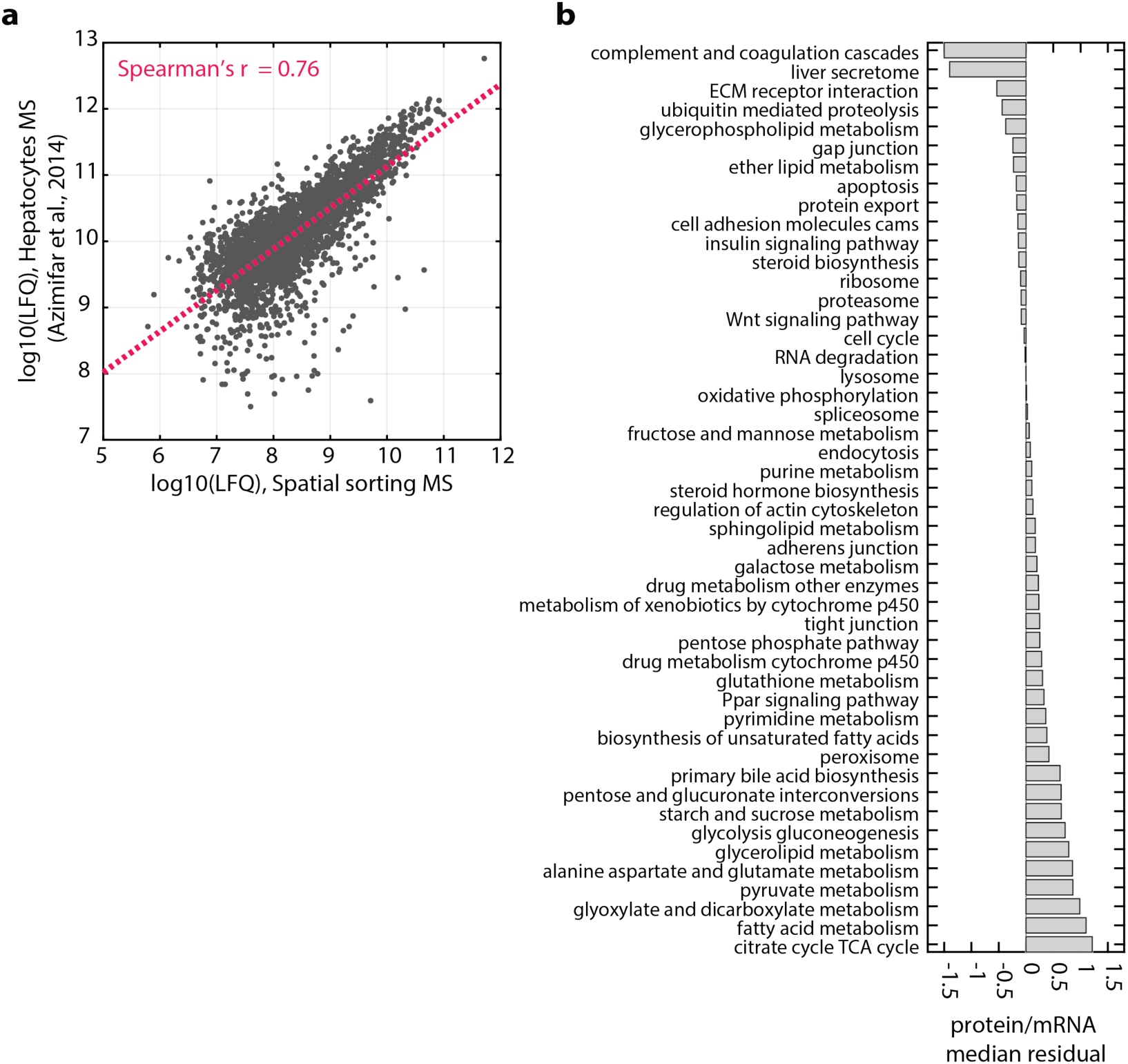
Comparisons of proteomic dataset. **a**, Scatter plot of log10(LFQ) of mean proteins over all FACS gates for 5 mice and log10(LFQ) of previously published mouse hepatocytes mass-spec. measurements ^12^. Red dashed line marks the regression line, Spearman’s r=0.76. Number of proteins = 2831. **b**, Protein to mRNA residuals from regression line, bar represent median of genes belonging to the KEGG pathway. See main Fig. 4.

**Supplementary Fig. 3.**
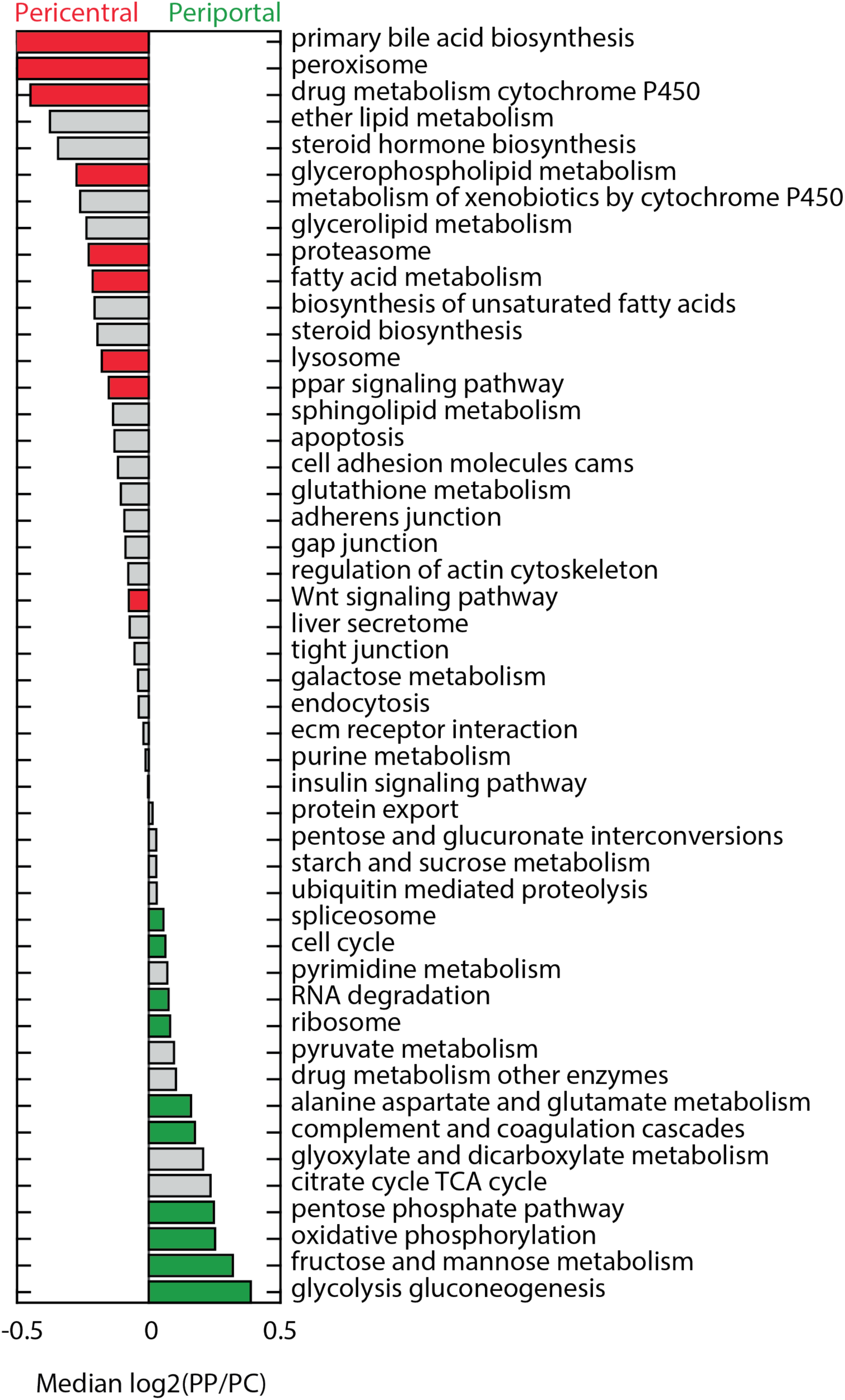
Zonation of hepatocyte proteome by KEGG pathways. Median log2 of ratio between periportal and pericentral gates of proteins belonging to the selected KEGG families. Colored bars represent KEGG pathways with FDR < 0.2 of single sample sign rank test p-values for each of the KEGG pathways. Red represent negative log2 ratio, indicating pericentral enrichment, and green represent positive values, indicating periportal enrichment.

**Supplementary Fig. 4.**
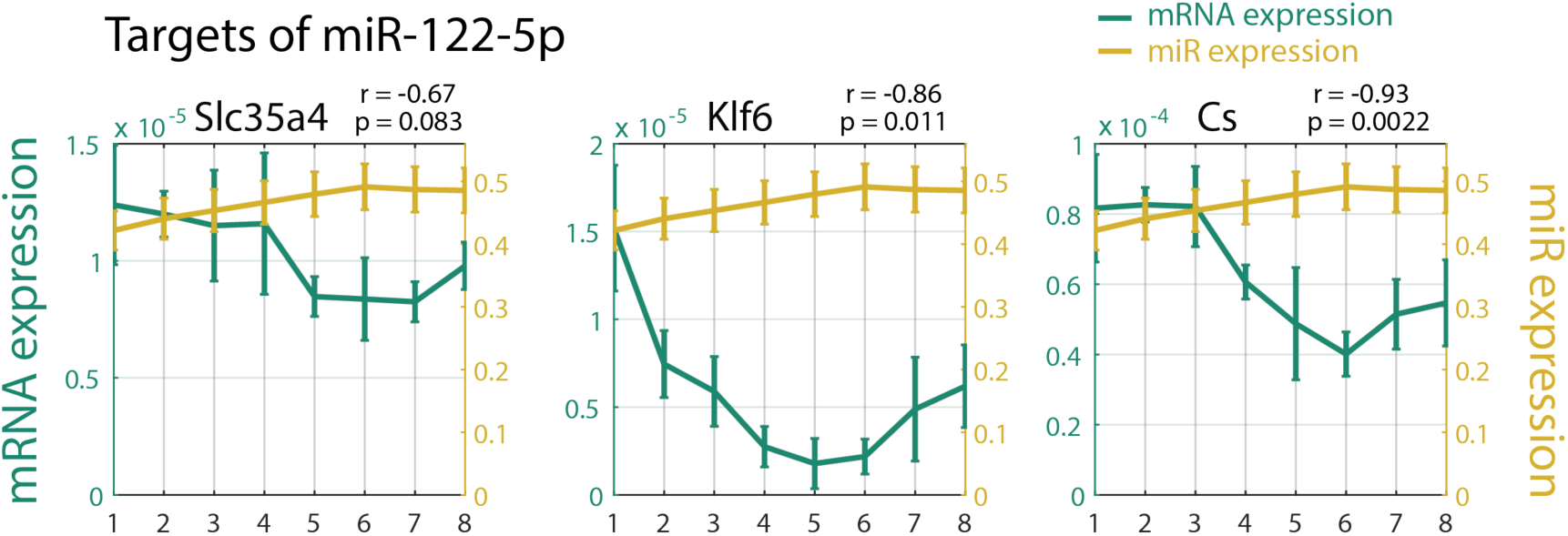
Selected anti-correlated target genes of miR-122-5p. MiR-122-5p was found to be anti-correlated with some of its canonical targets (following ^28,72^). Expression is normalized to the sum.

**Supplementary Fig. 5.**
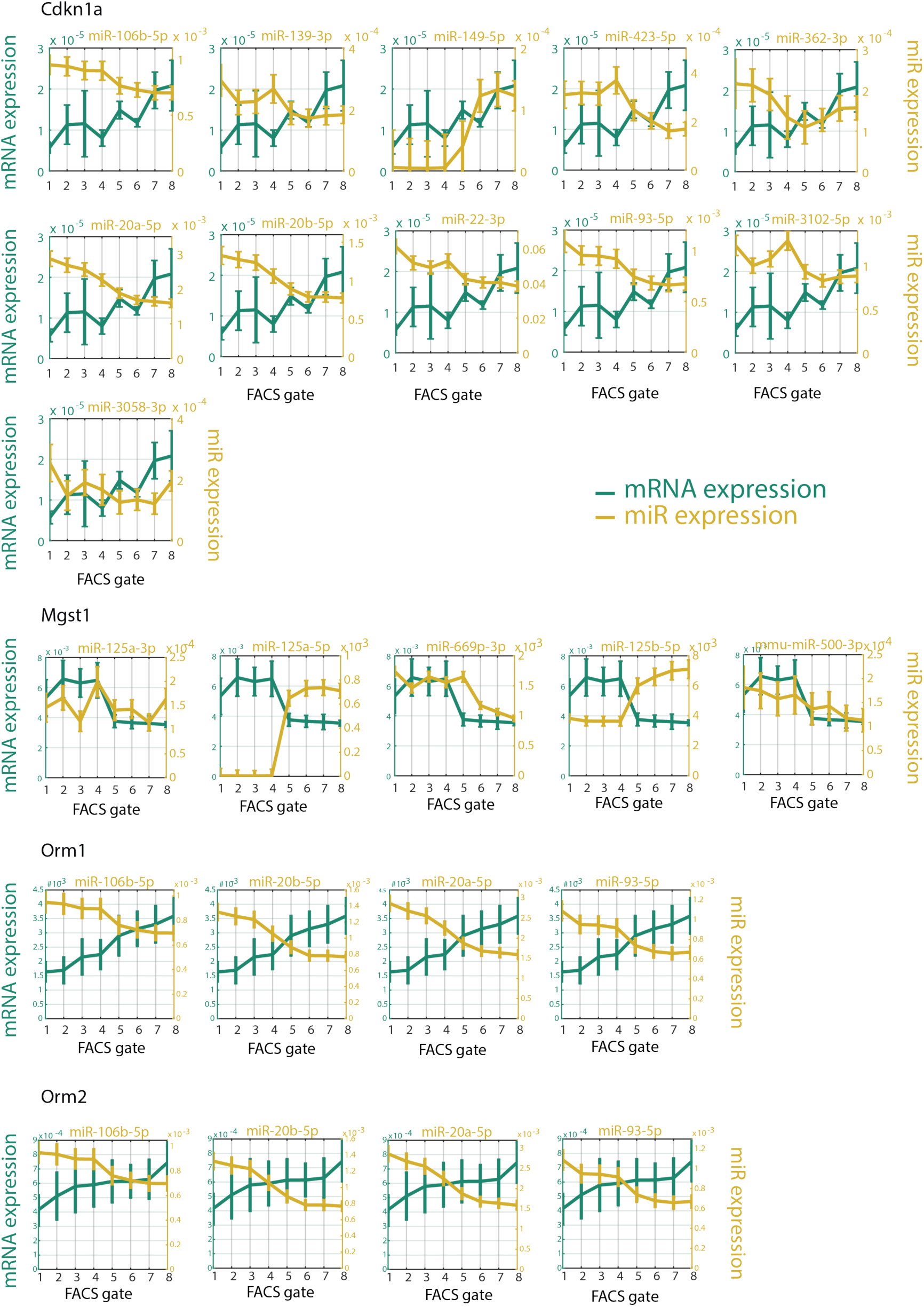
Individual expression profiles of select genes and their regulating miRs. Expression is normalized to the sum.

## Supplementary Tables

**Supplementary Table 1 - mRNA UMI counts**

Eight sorted populations (1-8) from five different mice (m1-m5) were collected and mRNA libraries were prepared according to mcSCRB-seq protocol ^62^, and zUMIs pipeline ^63^. Exonic UMI counts were normalized using TMM ^64^ implemented in EdgeR ^65^. The table values are the normalized CPM (UMI counts per million). Note that in our analyses, we have discarded m2_2 and m3_5 as their number of reads were less than 200K.

**Supplementary Table 2 - Proteins MS/MS quantification**

Eight sorted populations (1-8) from five different mice (m1-m5) were collected and extracted proteins were processed by LC-MS/MS. Data were analyzed and quantified with MaxQuant 1.5.2.8 ^66^ against the mouse Uniprot database. Table Values are log2(LFQ) – relative label-free quantification of proteins across samples. Table also shows sequence coverage, Razor+Unique peptides and MS/MS spectral count for each sample, as well as protein identifiers.

**Supplementary Table 3 - mRNA and proteins of spatially-sorted hepatocytes**

Shown are the data for 3,051 genes with matched transcriptomic and proteomic information. The mean and SEM of the five repeats in each FACS gate, normalized to their sample sum, were calculated for both mRNA and protein. Kruskal-Wallis p-value and FDR was assigned to each of the mRNAs/proteins.

**Supplementary Table 4 - Zonation profiles of detected high-confidence miRs**

Shown are the mean and SEM of the sum-normalized expression of miRs for each FACS gate, p-value for zonation (Kruskall-Wallis test), false discovery rate (FDR, using Benjamini-Hochberg procedure) and visualization of zonation profiles.

**Table 5. Zonation profiles of high-confidence miR-target pairs**

Includes expression level per FACS gate (as fraction of total) of miRs and their regulated genes, and Spearman correlation. Predicted interactions were obtained from TargetScan 7.2, with weighted context++ score percentile ≥ 95.

**Supplementary Table 6 - Zonation profiles of target genes and their cumulative miR profiles**

Includes expression level per FACS gate of genes and their cumulative miR profiles, Spearman correlation, p-value for the interaction as obtained by the network analysis (see Methods), FDR and visualization of gene and cumulative miR zonation profiles.

**Supplementary Table 7 - Primer sequences used for miR qRT-PCR**

